# Neonatal neuronal WWOX gene therapy rescues *Wwox* null phenotypes

**DOI:** 10.1101/2021.04.27.441575

**Authors:** Srinivasarao Repudi, Irina Kustanovich, Sara Abu-Swai, Shani Stern, Rami I. Aqeilan

## Abstract

WW domain-containing oxidoreductase (*WWOX*) is an emerging neural gene regulating homeostasis of the central nervous system. Germline biallelic mutations in *WWOX* cause WWOX-related epileptic encephalopathy (WOREE) syndrome and spinocerebellar ataxia, and autosomal recessive 12 (SCAR12), two devastating neurodevelopmental disorders with highly heterogenous clinical outcomes, the most common being severe epileptic encephalopathy and profound global developmental delay. We recently demonstrated that neuronal ablation of murine *Wwox* recapitulates phenotypes of *Wwox*-null mice leading to intractable epilepsy, hypomyelination and postnatal lethality. Here, we designed and produced an adeno-associated viral vector harboring murine *Wwox* or human *WWOX* cDNA and driven by the human neuronal Synapsin I promoter (*AAV-SynI-WWOX*). Testing the efficacy of AAV-SynI-WWOX delivery in *Wwox* null mice demonstrated that specific neuronal restoration of WWOX expression rescued brain hyperexcitability and seizures, hypoglycemia, and myelination deficits as well as the premature lethality of *Wwox*-null mice. These findings provide a proof-of-concept for *WWOX* gene therapy as a promising approach to curing children with WOREE and SCAR12.

## Introduction

The WW domain-containing oxidoreductase (*WWOX*) gene maps to chromosome 16q23.1-q23.2 encompassing the chromosomal fragile site FRA16D and encodes a 46kDa WWOX protein.^1,2^ WWOX comprises two WW domains (WW1 and WW2) and an extended short-chain dehydrogenase/reductase (SDR) domain.^3–5^ WWOX, via its WW1 domain, physically interacts with several key signaling proteins (Dvl, AP-2, ErbB-4, HIF1-α, p53, p63, p73, c-JUN, ITCH and RUNX2) and suppresses tumor progression in several cancer cell types.^6^ Additionally, WWOX has been shown to regulate DNA damage response, glucose homeostasis, cell metabolism and neuronal differentiation.^7,8^

In recent years, evidence linking WWOX function to the regulation of homeostasis of the central nervous system (CNS) has been proposed.^9,10^ Germline recessive mutations (missense, nonsense and partial/complete deletions) in the *WWOX* gene were found to be associated with two major phenotypes, namely SCAR12 (spinocerebellar ataxia, autosomal recessive -12, OMIM 614322) and WOREE syndrome (WWOX-related epileptic encephalopathy), the latter also known as developmental and epileptic encephalopathy-28 (DEE28, OMIM 616211).^9^ WOREE is a complex and devastating neurological disorder observed in children harboring an early premature stop codon or complete loss of *WWOX*.^11^ The clinical spectrum of WOREE includes severe developmental delay, early-onset of severe epilepsy with variable seizure manifestations (tonic, clonic, tonic–clonic, myoclonic, infantile spasms and absence). Most of the affected patients make no eye contact and are not able to sit, speak, or walk.^9^ WOREE syndrome is refractory to current anticonvulsant drugs, hence there is an urgent need to develop alternative treatments to help children with WOREE syndrome. Children with SCAR12, mostly due to missense mutations in *WWOX*, display a milder phenotype including ataxia and epilepsy.^12^ Epilepsy in SCAR12 can be treated with anticonvulsant drugs, though children still display ataxia and are intellectually disabled. Moreover, WWOX mutations have been documented in patients with West Syndrome, which is characterized by epileptic spasms with hypsarrhythmia.^13^ Brains of the children carrying *WWOX* gene mutations are found to be abnormal, as assessed by magnetic resonance imaging (MRI). Brain abnormalities such as hypoplasia of the corpus callosum, progressive cerebral atrophy, delayed myelination and optic nerve atrophy have been documented in most cases. It is largely unknown how mutations in *WWOX* or loss of WWOX function could lead to these CNS-associated abnormalities.

There is a marked similarity between human WWOX (*hWWOX*) and murine Wwox (*mWwox*). In fact, the human WWOX protein sequence is 93% identical and 95% similar to the murine WWOX protein sequence. Remarkably, targeted loss of *Wwox* function in rodent models (mice and rats) phenocopies the complex human neurological phenotypes, including severe epileptic seizures, growth retardation, ataxia and premature death.^12,14,15^ *Wwox* null mice also exhibit phenotypes associated with impaired bone metabolism and steroidogenesis.^16,17^ In a recent study, Repudi et al. found that conditional ablation of murine *Wwox* in either neural stem cells and progenitors (N-KO) or neuronal cells (S-KO mice) resulted in severe epilepsy, ataxia and premature death at 3-4 weeks, recapitulating the phenotypes observed in the *Wwox*-null mice.^18^ These results highlight the significant role of WWOX in neuronal function and prompted us to test whether restoring WWOX expression in the neuronal compartment of *Wwox* null mice could reverse the observed phenotypes. To this end, we used an adeno-associated virus (AAV) vector to restore WWOX expression. AAV is a promising candidate for gene therapy in many disorders including neuromuscular, CNS and ocular disorders.^19–22^ Moreover, AAV appears to elicit little to no immune response and integrates into the host at very low rates, which reduces the risks of genotoxicity.^23^ In our study, we demonstrated that an AAV vector harboring the *mWwox* or *hWWOX* open reading frame and driven by the human neuronal Synapsin I promoter could reverse *Wwox* null phenotypes. A single intracerebroventricular (ICV) injection of *AAV9-Synapsin I-WWOX* rescued the growth retardation, epileptic seizures, ataxia and premature death of *Wwox* null mice. In addition, WWOX restoration improved myelination and reversed the abnormal behavioral changes of *Wwox* null mice. Overall, these remarkable results indicate that WWOX gene therapy could be a promising cure approach for children with WOREE and SCAR12.

## Results

### Restoration of neuronal WWOX rescues growth retardation and post-natal lethality of *Wwox* null mutant mice

In a recent study, we reported that conditional ablation of WWOX in neurons phenocopies the *Wwox* null mice including growth retardation, spontaneous epileptic seizures, ataxia and premature death at 3-4 weeks.^18^ These results implied that WWOX is a key neuronal gene regulating homeostasis of the CNS. Prompted by these remarkable findings, we wanted to address whether neuronal-specific expression of WWOX in *Wwox*-null mice could rescue lethality of these mice and their associated phenotypes. We designed an adeno-associated viral (AAV) vector to express murine *Wwox (mWwox),* or human *WWOX* (*hWWOX*), cDNA driven by a human Synapsin-I (hSynI) promoter (**Fig. 1A**) and packaged this into an AAV9 serotype, which has high CNS tropism and has been used in CNS-based gene therapy trials.^22,24^ An *IRES-EGFP* sequence was cloned downstream of the *Wwox/WWOX* sequence to allow tracking of expression. Successful delivery of AAV9-hSynI-*mWwox*-IRES-EGFP (*AAV9-hSynI-mWwox*) should lead to expression of intact WWOX protein in Synapsin-I-positive non-dividing/matured neurons. As control, *AAV9-hSynI-EGFP* was used. Expression of WWOX and GFP was initially validated by infecting primary *Wwox*-null dorsal root ganglion (DRGs) neurons with the viral particles *in vitro* (**Supplementary Fig. S1**).

**Figure 1.**
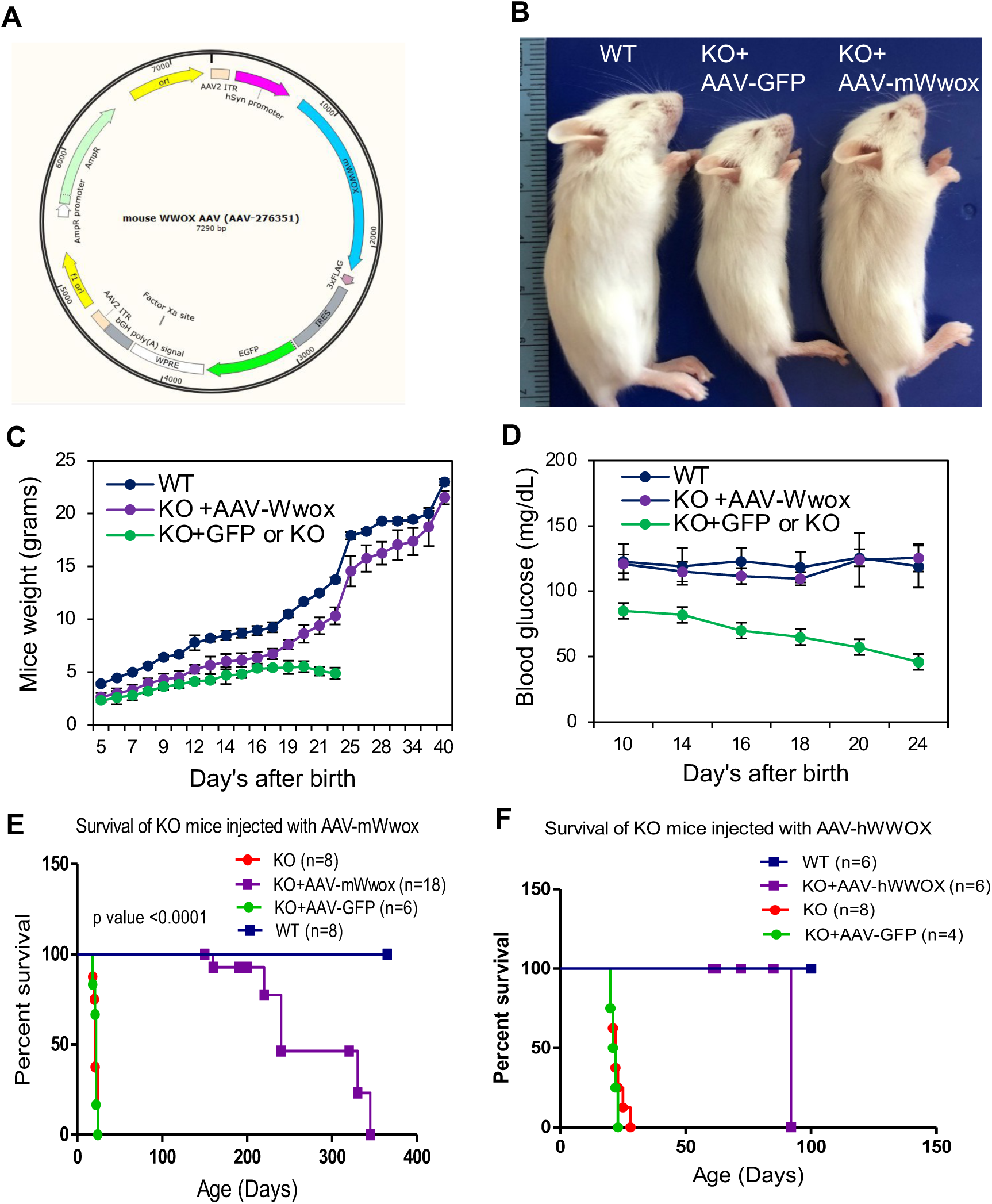
Restoration of WWOX in Synapsin I-positive neurons improves growth of *Wwox* null mice and extends their life span. (**A**) Illustration of the plasmid vector construct containing murine *Wwox* gene (shown in blue) under human Synapsin I promoter. The *Wwox* gene sequence is followed by an IRES promoter and *EGFP* gene sequence. (**B**) Physical appearance of wild type (WT), *Wwox* null injected either with *AAV9-hSynI-GFP* (the control virus) or *AAV9-hSynI-mWwox-IRES-GFP* virus at P17. Graphs showing body weight (**C**) and blood glucose levels (**D**) of the mice at indicated days. Error bars represent ±SEM, n=4 mice per genotype. (**E**) Kaplan-Meier survival graph indicates prolonged life span of *Wwox* knockout mice injected with *AAV9-hSynI-mWwox* (n=18) compared to mice injected with *AAV9-hSynI-GFP* (n=6) or the non-injected (n=10) (p value <0.0001, Log-rank Mantel-Cox test). (**F**) Kaplan-Meier survival graph indicates prolonged life span of *Wwox* knockout mice injected with *AAV9-hSynI-hWWOX* (n=6) [median 92 days] compared to mice injected with *AAV9-hSynI-GFP* (n=4) or the non-injected (n=8) (p value = 0.0001, Log-rank Mantel-Cox test).

We then evaluated the expression and function of the AAVs *in vivo*. Viral particles (2×10^10^/hemisphere) of *AAV9-hSynI-mWwox* or *AAV9-hSynI-EGFP* were injected into the intracerebroventricular region of *Wwox* null mice at birth (P0), to achieve widespread transduction of neurons throughout the brain.^25,26^ Successful expression of the transgene in neurons, but not in oligodendrocytes (CC1-positive cells), was validated by immunofluorescence using anti-NeuN and anti-WWOX antibodies (Fig. 2A, B and C).

**Figure 2.**
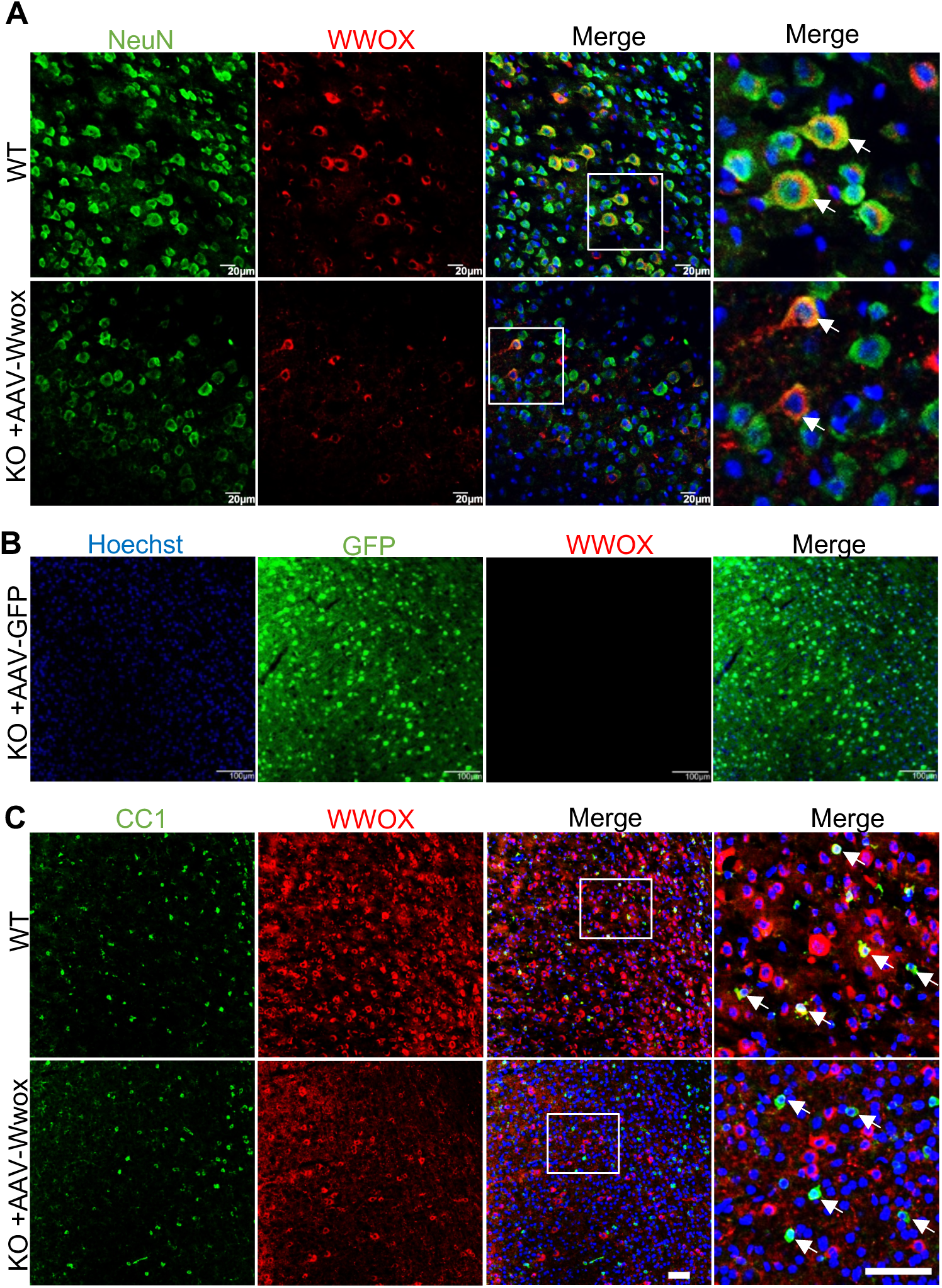
Validating expression of the transgene (WWOX or GFP) in AAV injected mice brain. (**A**) Immunofluorescence images showing the expression of WWOX (red) in brain tissue (cortex) at P17 from WT and *Wwox* null mice that were injected with *AAV9-hSynI-mWwox* (2×10^10^). Neurons are labelled with anti-NeuN antibody (green). The square white box shows the magnified area. (**B**) Expression of EGFP (green) is shown in brain tissue (cortex) of *Wwox* null injected with *AAV9-hSynI-GFP* (2×10^10^). (**C**) Brain sections were immunolabelled with CC1 and anti-WWOX showing WWOX (red) expression in oligodendrocytes (shown with arrow in WT panel) of WT and not in *Wwox* null (shown with arrow) injected with *AAV9-hSynI-mWwox*. Representative images are shown from the cortex region. Scale bars **A**) 20 µm, **B**) 100 µm **C**) 20 µm.

Monitoring of the treated mice revealed that mice injected with *AAV9-hSynI-mWwox* grew normally (**Fig. 1B**), gradually gained weight (**Fig. 1C**) and were indistinguishable from wild type by the age of 6-8 weeks. *AAV9-hSynI-EGFP*-injected mice exhibited similar phenotypes of *Wwox* null mice (**Fig. 1C**). Of note, *Wwox* null and *AAV9-hSynI-EGFP*-injected mice were hypoglycemic from the second week until they died, while *AAV9-hSynI-mWwox*-injected mice had normal blood glucose levels when compared to the wild type mice (**Fig. 1D**). Remarkably, all rescued mice lived longer with a median survival of 240 days compared to the *Wwox* null or *AAV9-hSynI-EGFP*-injected *Wwox* null mice (p value <0.0001) (**Fig. 1E**). Similar results and outcomes were obtained when replacing *mWwox* with *hWWOX* cDNA, though we have only followed these mice for up to 100 days so far (**Fig. 1F**, **Supplementary Fig. S2**). Notably, the rescued mice were active and both males and females were fertile (data not shown). Since *Wwox* null mice were previously shown to lack testicular Leydig cells,^16^ we next determined if WWOX neuronal restoration rescues this phenotype and indeed found intact Leydig cells in P17 *AAV9-hSynI-mWwox*-treated mice (**Supplementary Fig. S3A**). Bone growth defects were also formerly documented in *Wwox* mutant mice^17,27–30^, and hence we examined bones of rescued mice and observed that cortical bones were of comparable size and thickness to WT mice (**Supplementary Fig. S3B**). These results imply that neuronal restoration of WWOX could be sufficient to rescue the abnormal phenotypes of *Wwox* null mice.

### Neuronal restoration of WWOX decreases hyperexcitability of *Wwox* null mice

We and others have previously reported that *Wwox* null mutants display spontaneous recurrent seizures.^12,14,18,31^ As we did not observe any spontaneous seizures in rescued mice, we determined next the epileptic activity in brains of P21-22 wild type (WT), *Wwox* null (KO) and *AAV9-hSynI-mWwox*-injected *Wwox* null mice (KO*+AAV9-Wwox*) by performing cell-attached electrophysiology recordings (**Supplementary Fig. S3**). As expected, the KO pups exhibited severe hyperactivity. Representative traces with spontaneous firing of action potentials are shown in **Fig. 3A**. A clear hyperexcitability can be noted from the representative traces (WT in blue, KO+*AAV9-Wwox* in purple and KO in red). The activity of the KO brains usually resulted in bursts of action potentials and overall there was a drastic increase in the firing rate. The average firing rate over 20 WT, 20 KO*+AAV-Wwox* and 30 KO recorded neurons was about 6-fold higher in KO pups compared to the WT pups (p=2.6e-7). No significant difference in average firing rate was observed between the KO+*AAV-Wwox* and the WT pups.

**Figure 3.**
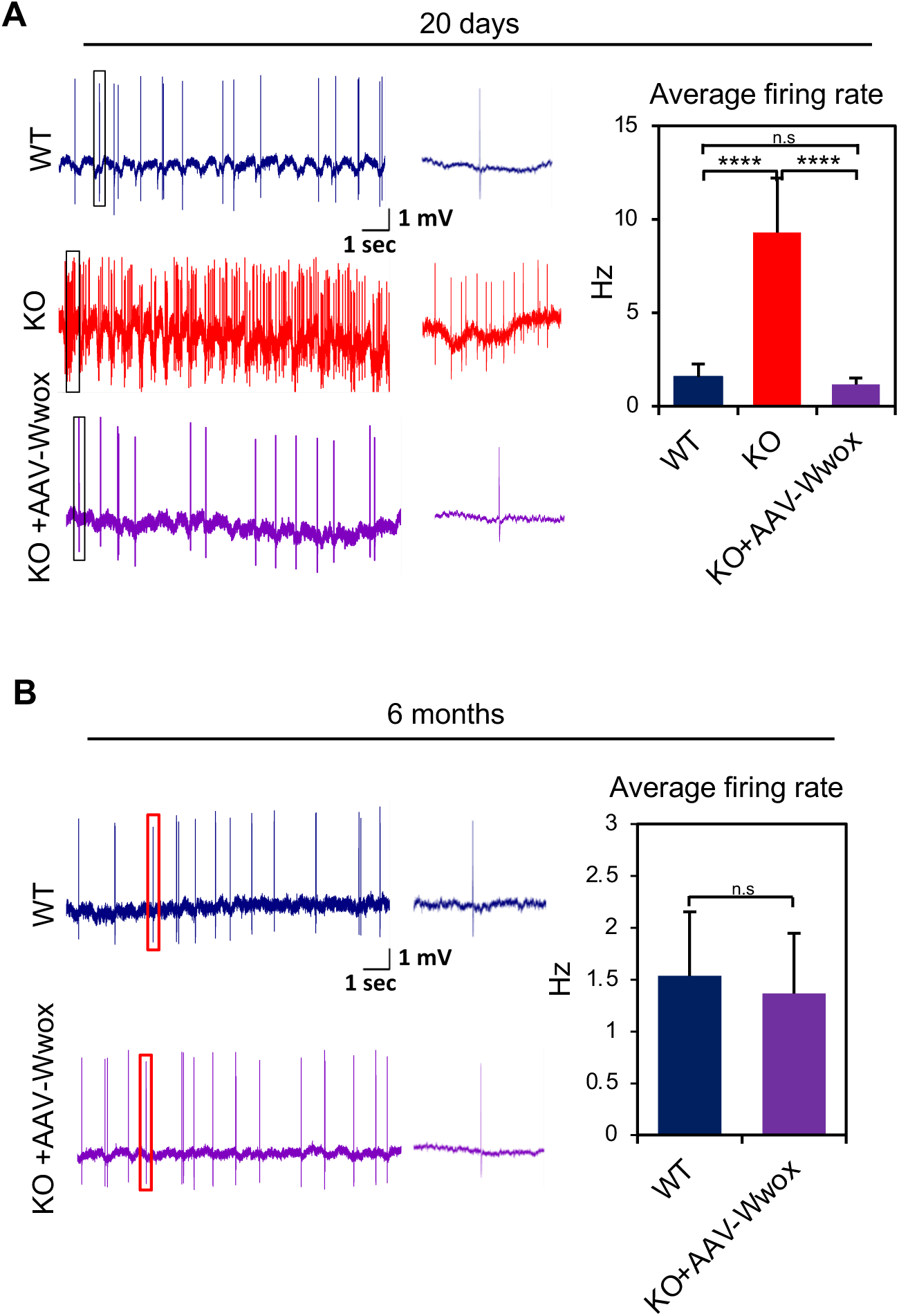
Neuronal restoration of WWOX reduces epileptic activity in neocortex. (**A**) Representative traces of cell-attached recordings performed in WT (blue), KO (red) and KO-treated with *AAV9-hSynI-mWwox* [KO+A-Wwox] (purple) pups at P20-21 days. Traces represent spontaneous neocortical activity (action potentials shown). The panels represent 12 s recording with an inset presenting a zoom-in of an 0.5 s interval. A clear hyperactivity of the KO brain is observed in these representative traces with the KO showing bursts of action potentials. The graph shows the averages over 20 neurons from WT pups (n=2), 20 neurons from KO+A-Wwox pups (n=2) and 30 neurons from KO pups (n=2) (****p value <0.0001, student’s t test). (**B**) Representative traces of cell-attached recordings performed in a WT (blue) and a KO+A-Wwox (purple) adult mice (6 months old). Traces present spontaneous neocortical activity (action potentials). The panels present 12 s recording with an inset presenting a zoom-in of an 0.5 s interval. The graph shows the averages over 60 neurons from WT adult mice (n=3) and 60 neurons from KO+A-Wwox adult mice (n=3). There was no significant change observed between the average firing rate of the WT and the KO+A-Wwox adult mice.

Since KO mice died within less than 4 weeks, we could not perform *in vivo* recordings in adult KO mice. We therefore performed cell attached *in vivo* recordings only in adult WT and KO+*AAV9-Wwox* mice (**Fig. 3B**). Representative traces are shown in blue (WT) and purple (KO+*AAV9-mWwox*) and the average firing rate over 60 WT and 60 KO+*AAV9-mWwox* neurons are presented (**Fig. 3B**). There were no significant differences in the firing rate of adult WT and KO+*AAV9-mWwox* cortical neurons. These findings indicate that *AAV9-hSynI-mWwox* could prevent epileptic seizures resulting from WWOX loss.

### Neuronal restoration of WWOX enhances myelination in *Wwox* null mice likely by promoting OPC differentiation

Previous observations linked WWOX loss with hypomyelination.^18,31^ In fact, it was shown that neuronal WWOX ablation results in a non-cell autonomous function impairing differentiation of oligodendrocyte progenitors (OPCs).^18^ Hence we next tested whether neuronal restoration of WWOX, using AAV, could rescue the hypomyelination phenotype in *Wwox* null mice. Immunofluorescence analysis of P17 sagittal brain tissues with anti-MBP antibody revealed improved myelination in all parts (cortex, hippocampus and cerebellum) of the rescued *AAV9-hSynI-mWwox* treated-mice brain compared to *Wwox* null mice injected with control virus (**Fig. 4A**). In addition, we tested whether this improved myelination is associated with increased differentiation of OPCs to matured oligodendrocytes by a non-cell autonomous function of neuronal WWOX. As expected, AAV9-mediated WWOX expression in neurons increased the differentiation of OPCs to matured oligodendrocytes as assessed by immunostaining with CC1 (marker for matured oligodendrocytes) and anti-PDGFRα (marker for OPCs) (**Fig. 4B**). Quantification of CC1 and OPCs in the corpus callosum showed significantly increased number of matured oligodendrocytes in rescued mice compared to the KO mice injected with control virus on P17 (**Fig. 4C**).

**Figure 4.**
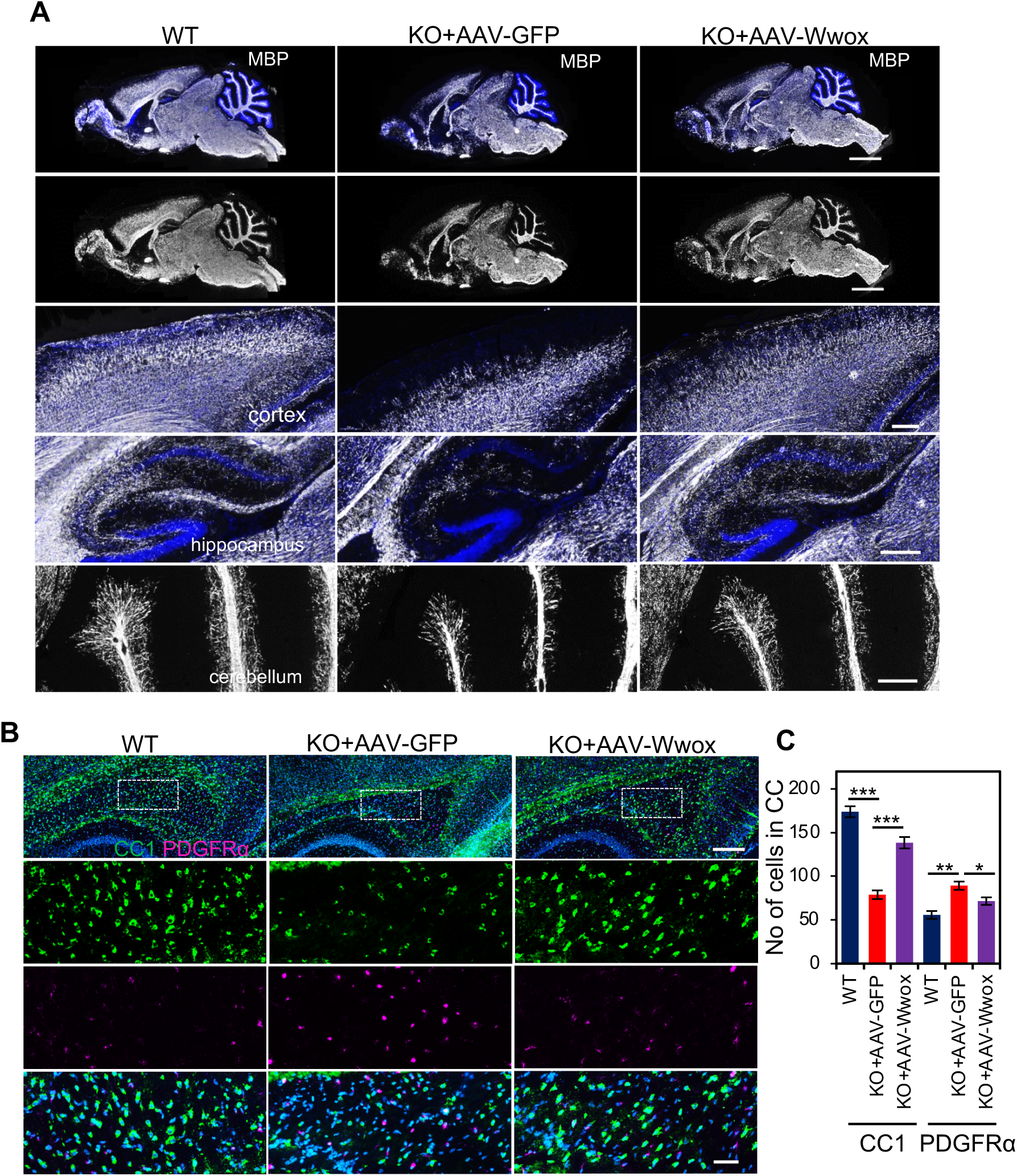
WWOX restoration in neurons improves myelination by promoting OPCs differentiation in *Wwox* null. (**A**) Images of whole brain sagittal sections that were immunolabelled with anti-MBP at P17 from indicated mice (n=3 for each group). MBP staining (shown in gray) in cortex, hippocampus and cerebellum is presented (magnified from the top panel). (**B**) Sagittal section of the brain (P17) tissues immunostained for CC1 (green) and PDGFRα (magenta). Images showing increased number of matured oligodendrocytes in corpus callosum of the *Wwox* null after treatment with *AAV9-hSynI-mWwox* compared to *Wwox* null injected with *AAV9-hSynI-GFP* virus. (**C**) Graph represents the quantification of CC1 and PDGFRα positive cells in corpus callosum (area 0.5 mm^2^) of WT (n=3), KO (n=3) and KO+A-Wwox (n=3) counted from three similar brain sagittal sections per mouse in each genotype. Error bars represent ±SEM. (*p value <0.01, **p value <0.001, ***p value <0.0001). Scale bars **A**) 2 mm (top panel), 250 µm (middle and lower panels), **B**) 250 µm (top panel), 50 µm (middle and lower panels).

To further validate the finding of improved myelination after neuronal restoration of WWOX, we performed electron microscopy (EM) analysis for corpus callosum on P17 and in adult mice. Remarkably, neuronal restoration of WWOX using *AAV9-hSynI-mWwox* increased the number of myelinated axons compared to KO at P17 (**Fig. 5A, B**) in the corpus callosum. Furthermore, calculated g-ratios indicated increased myelin thickness upon neuronal WWOX restoration compared to control KO mice (**Fig. 5C**). In addition, EM images of the corpus callosum and optic nerves of adult (6 months) rescued mice showed improved myelination (**Fig. 5D-F, data not shown**). Of note, when comparing myelin thickness of KO+*AAV-Wwox* and WT corpus callosum at P17 and 6 months, we observed some differences in g-ratio (**Fig. 5C and F**).

**Figure 5.**
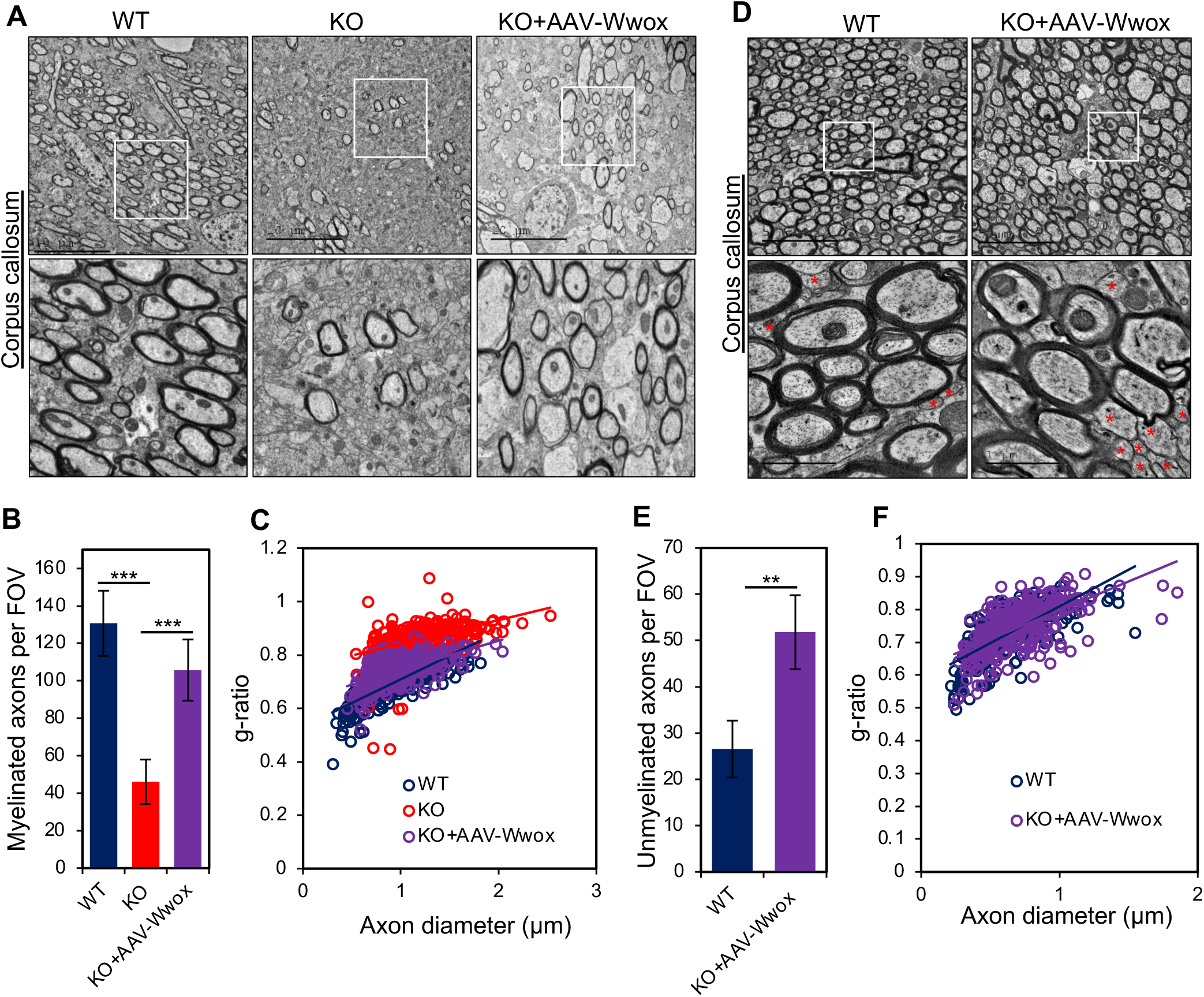
Electron microscopy analysis showing improved myelination in corpus callosum upon WWOX restoration. (**A**) Electron micrograph (EM) images from mid sagittal section of corpus callosum showing increased number of myelinated axons in *AAV9-hSynI-mWwox*-injected mice compared to *Wwox*-null mice (KO) at P17. (**B**) Graph showing average number of myelinated axons in corpus callosum from WT (n=3), KO (n=3) or KO mice injected with *AAV9-hSynI-mWwox* (n=3) per field of view (FOV). (***p value <0.0001). (**C**) g-ratio analysis showing reduced myelin thickness of axons in corpus callosum KO (n=3) compared to the KO+AAV-Wwox (n=3). Axon diameter and myelin thickness was calculated from the electron micrograph images of corpus callosum (n=300 per each genotype). (**D**) EM images from mid sagittal section of corpus callosum showing similar number of myelinated axons in rescued mice (*AAV9-hSynI-mWwox*, n=2) compared to wild type (n=2) at the age of 6 months. Unmyelinated axons are noted with the red star symbols in EM images. (**E**) Graph represents the number of unmyelinated axons (corpus callosum) per FOV (**p value <0.001). (**E**) Graph represents the measured g-ratio of myelinated axons (n=300 per each genotype) in WT and rescued mice at 6 months (p value 0.002334). Scale bars **A**) 10 µm (top panel), **D**) 5 µm (inset 1 µm).

### WWOX neuronal restoration decreases anxiety and improves motor functions

We next explored the behavioral changes in *Wwox* null mice after restoration of WWOX in neurons. Unfortunately, we could not assess behavior of *Wwox* null mice due to their poor conditions and premature death. We performed open field, elevated plus maze (EPM) and rotarod tests to examine anxiety and motor coordination in rescued mice (**Fig. 6**). Remarkably, at 8-10 weeks we observed similar tracking patterns in the open field in the rescued mice (males and females) to those seen in the wild type, indicating that WWOX restoration reduces anxiety in *Wwox* null mice. In addition, the velocity and total distance travelled in the open field tracks were very similar to that of the WT mice (**Fig. 6B, C**). Moreover, rescued female and male mice exhibited near normal behavior in the EPM (**Fig. 6D**). Rotarod test was performed to check the motor coordination in rescued mice. Our results revealed that rescued mice had similar motor coordination to WT mice in trial 3, an indicative of their learning ability. Altogether, these results suggest that neuronal WWOX re-expression in *Wwox* null mice restores activity and normal behavior.

**Figure 6.**
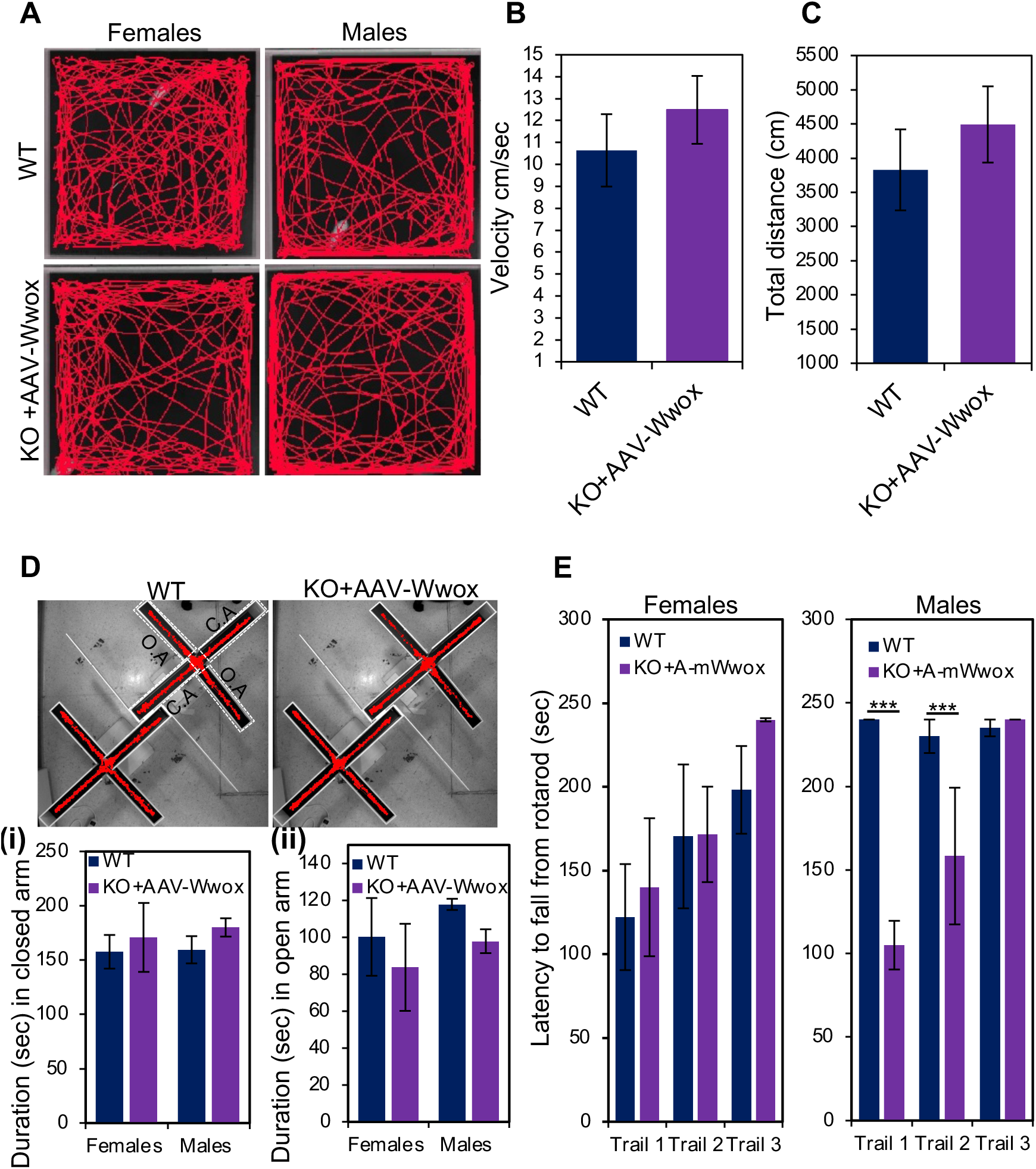
WWOX restoration in neurons reverses the abnormal behavioral phenotypes of *Wwox* null mice. (**A**) Representative images of open field test showing tracking pattern (periphery and the center) of WT (females, n=7, males n=6) and *AAVmWwox*-injected KO mice (females n=7, males n=5) at 8-10 weeks. (**B**) and (**C**) Graphs represent the velocity and distance travelled (cm) from the open field tracking. (**D**) Mice tracking images of elevated plus maze test from WT and KO+AAV-Wwox mice (age 8-10 weeks). O. A. (open arm) C.O. (closed arm) are shown with dotted white boxes on the left mice tracking image. Graphs represent the duration (s) in closed arm (i) and open arm (ii) from WT (females, n=7, males n=6) and KO+AAV-Wwox (females n=7, males n=5). (**E**) Graphs represent the latency (s) of the mice (WT and rescued mice at 8-10 weeks) to fall from the rotarod in different trials. Females (WT, n=7; KO+*-*mWwox, n=5) and males (WT, n=5; KO+*-*mWwox, n=5) are shown separately (***p value <0.0001).

## Discussion

Based on our recent findings^18^ we aimed here to restore WWOX in neurons and assess the therapeutic potential of this restoration. In this study, we utilized an AAV9 vector for targeted gene delivery of WWOX to mature neurons to treat the complex neuropathy in the *Wwox*-null mouse model. We injected *mWwox* or *hWWOX cDNA* under the neuronal promotor *Synapsin-I* into the brains of newborn *Wwox* null mice and showed that this treatment was able to reverse the phenotypes of WWOX deficiency.

The role of WWOX in regulating CNS homeostasis is emerging as a key function of the *WWOX* gene. Deficiency of WWOX has been linked to a number of neurological disorders.^9,10^ Of particular interest is WOREE syndrome, a devastating complex neurological disease causing premature death with a median survival of 1-4 years.^9,10^ WOREE children are refractory to the current antiepileptic drugs (AEDs) hence challenging the medical and scientific communities to develop new therapeutic strategies. We believe that delivering AAV9-WWOX into the brain of WOREE syndrome patients could be a novel gene therapy approach that would help these patients. Recent success in gene therapy clinical trials of the treatment of spinal muscular atrophy (SMA) using the AAV9 vector^32^ is encouraging and has promoted our further development of this platform in this proof-of-concept study.

The effects of delivering *AAV9-SynI-WWOX* into the brains of *Wwox* null mice were remarkable. Firstly, WWOX neuronal delivery restored normal growth and survival of mice with no occurrence of spontaneous seizures and ataxia. In addition, we showed that neuronal restoration of WWOX reduced hyperexcitability in cell-attached recordings. Secondly, neuronal WWOX restoration improved myelination of all regions of the brain further confirming the previous observations of WWOX neuronal non-cell autonomous function on OPC maturation.^18^ Of note, there are still some differences between rescued and WT mice which could be attributed to an oligodendrocyte-specific WWOX function in regulating the myelination process. Thirdly, WWOX restoration improved the overall behavior of the rescued mice. These findings might suggest that WWOX’s proposed role in regulating autism^9–11,33,34^ and perhaps other behavior-associated disorders is driven by proper neuronal function of WWOX.

Another intriguing consequence of neuronal WWOX delivery is the reversibility of hypoglycemia associated with WWOX deficiency in *Wwox* null mice.^35,36^ These results are consistent with a central role of WWOX in the CNS regulation of metabolism of glucose and likely other metabolic functions.^37–40^ Interestingly, targeted deletion of *Wwox* in skeletal muscle resulted in impaired glucose homeostasis^41^ and this effect was linked to cell-autonomous functions of WWOX. Another peculiar observation is that the rescued mice were also fertile and able to breed. Given that *Wwox* null mice were shown to display impaired steroidogenesis^16,29,42^, our current findings imply that WWOX’s function in the CNS is superimposing its tissue level function. Altogether, these findings suggest that WWOX could have pleotropic function both at the organ level and at the organism level.

WWOX is ubiquitously expressed in all brain regions.^10,43,44^ Our current observations do not imply that WWOX expression in other brain cell types, such as astrocytes and oligodendrocyte, are dispensable. Evidence linking WWOX function with oligodendrocyte pathology is starting to emerge^45–49^, however less is known about the cell-autonomous functions of WWOX in oligodendrocytes. The fact that WWOX expression in neurons regulates oligodendrocyte maturation and antagonizes astrogliosis^50^ suggests a complex function of WWOX in CNS physiology and pathophysiology that warrants further in-depth analysis.

The *WWOX* gene was initially cloned as a putative tumor suppressor.^51,52^ Indeed a plethora of research work in various animal models (reviewed in^15^) and observations in human cancer patients^1,27,39,53–57^ proposed WWOX as a tumor suppressor. Given that our restoration of WWOX is limited to brain, we assumed other tissues lacking WWOX expression would be more susceptible to tumor development. Of note, we didn’t detect gross tumor formation in the limited number of adult *Wwox*-null mice treated with *AAV9-hSynI-mWwox* that we examined (age 6-8month, n=6). This was not surprising given that *Wwox* somatic deletion in several tissues required other hits to promote tumor formation in animal models.^28,39,58,59^ Nevertheless, detailed cellular and molecular analyses shall be required in the future to further investigate any abnormal changes associated with WWOX deficiency in AAV-treated mice.

The limited life-span and poor conditions of *Wwox* null mice forced us to treat these mice very early on in their life (P0). Nevertheless, attempts to treat post-natal *Wwox*-null mice should be explored in the future. Our current findings indicate that WWOX restoration in neonatal mice using an AAV vector could reverse the phenotypes associated with WWOX deficiency. We envisage that this proof-of-concept will lay down the groundwork for a possible gene therapy clinical trial on children suffering from the devastating and often refractory WOREE syndrome.

## Materials and Methods

### Plasmid vectors

Murine *Wwox* or human *WWOX* cDNA was cloned under the promoter of human *Synapsin I* in pAAV and this vector was packaged into AAV9 serotype (Vector Biolabs, Philadelphia, USA). Custom-made *AAV9-hSynI-mWwox-IRES-EGFP*, *AAV9-hSynI-hWWOX-2A-EGFP* and *AAV9-hSynI-EGFP* viral particles were obtained either from Vector Biolabs or from the Vector Core Facility at Hebrew University of Jerusalem.

### Mice

Generation of *Wwox* null ^(-/-)^ mice (KO) was previously reported^16^ and these mice were maintained in an FVB background. Heterozygote ^(+/-)^ mice were used for breeding to get the *Wwox* null mice. Animals were maintained in a SPF unit in a 12 h-light/dark cycle with *ad libitum* access to the food and water. All animal-related experiments were performed in accordance and with prior approval of the Hebrew University-Institutional Animal Care Use Committee (HU-IACUC).

### Intracerebroventricular (ICV) injection of AAV particles in to P0 *Wwox* null mice

Free-hand intracranial injections of either *AAV9-hSynI-mWwox-IRES-EGFP (AAV9-WWOX)* or *AAV9-hSynI-EGFP (AAV9-GFP)* into the *Wwox* null mice were done following a published protocol.^25^ Briefly, when neonates were born, they were PCR genotyped to identify *Wwox* null mice. *Wwox* null neonates were anesthetized by placing on a dry, flat, cold surface. The anesthetized pup head was gently wiped with a cotton swab soaked in 70% ethanol. Trypan blue 0.1% was added to the virus to enable visualization of the dispensed liquid. An injection site was located at 2/5 of the distance from the lambda suture to each eye. Holding the syringe (preloaded with virus) perpendicular to the surface of the skull, the needle was inserted to a depth of approximately 3 mm. Approximately 1 µl (2 x 10^10^ GC/hemisphere) virus was dispensed using a NanoFil syringe with a 33G beveled needle (World Precision Instruments). The other hemisphere was injected in the same way. Injected pups were placed on the warming pad until they were awake, then transferred to the mother’s cage. Each injected mouse was carefully monitored for growth, mobility, seizures, ataxia and general condition to assess phenotypes.

### Weight and blood glucose levels

Mice were weighed regularly as indicated in the Figures. To monitor the blood glucose, the tip of the mouse tail was ruptured with scissors and a tiny drop of blood collected for measurement (mg/dL) using an Accu-Check glucometer (Roche Diagnostics, Mannheim, Germany).

### Immunofluorescence

Mice from different genotypes and treatment groups (P17-P18) were euthanized by CO_2_ and transcardially perfused with 2% PFA/PBS. Dissected brains were postfixed on ice for 30 min then incubated in 30% sucrose at 4°C overnight. They were then embedded in OCT and sectioned (12-14 µm) using a cryostat. Sagittal sections were washed with PBS and blocked with 5% goat serum containing 0.5% Triton X-100 then incubated for 1 h at room temperature followed by incubation with primary antibodies overnight at 4°C. Then, sections were washed with PBS and incubated with corresponding secondary antibodies tagged with Alexa fluorophore for 1 h at room temperature followed by washing with PBS and mounting with mounting medium.

### Surgical procedures for electrophysiology

Mice were anesthetized using ketamine/medetomidine (i.p; 100 and 83 mg/kg, respectively). The effectiveness of anesthesia was confirmed by the absence of toe-pinch reflexes. Supplemental doses were administered every ∼1 h with a quarter of the initial dosage to maintain anesthesia during the electrophysiology procedures. During all surgeries and experiments, body temperature was maintained using a heating pad (37°C). The skin was removed to expose the skull. A custom-made metal pin was affixed to the skull using dental cement and connected to a custom stage. A small hole (3 mm diameter craniotomy) was made in the skull using a biopsy punch (Miltex, PA).

### Cell attached recordings

Cell-attached recordings were obtained with blind patch-clamp recording. Electrodes (∼7 MOhm) were pulled from filamented, thin-walled, borosilicate glass (outer diameter, 1.5 mm; inner diameter, 0.86 mm; Hilgenberg GmbH, Malsfeld, Germany) on a vertical two-stage puller (PC-12, Narishige, EastMeadow, NY). The electrodes were filled with internal solution containing the following: 140 mM K-gluconate, 10 mM KCl, 10 mM HEPES, 10 mM Na_2_-phosphocreatine, and 0.5 mM EGTA, adjusted to pH 7.25 with KOH. The electrode was inserted at a 45 degrees angle and reached a depth of 300 µm. The electrode positioning was targeted on the brain surface, positioned at 1.6-2 mm posterior to the bregma and 4 mm lateral to the midline. While positioning the electrode, an increase of the pipette resistance to 10–200 MOhm resulted in most cases in the appearance of action potentials (spikes). The detection of a single spike was the criteria to start the recording. All recordings were acquired with an intracellular amplifier in current clamp mode (Multiclamp 700B, Molecular Devices), acquired at 10 kHz (CED Micro 1401-3, Cambridge Electronic Design Limited) and filtered with a high pass filter. For calculation of the average firing rate, the firing rate over a 4 min recording period was calculated for each of the recorded cells. A two sample t-test was used to assess statistical significance between the recorded groups.

### Electron microscopy

Mice were anesthetized and perfused with a fixative containing 2% paraformaldehyde and 2.5% glutaraldehyde (EM grade) in 0.1 M sodium cacodylate buffer, pH 7.3. Brains were isolated and incubated in the same fixative for 2 h at room temperature then stored in 4°C until they were processed. Collected tissues (corpus callosum, optic nerve) were washed four times with sodium cacodylate and postfixed for 1 h with 1% osmium tetroxide, 1.5% potassium ferricyanide in sodium cacodylate, and washed four times with the same buffer. Then, tissue samples were dehydrated with graded series of ethanol solutions (30, 50, 70, 80, 90, 95%) for 10 min each and then 100% ethanol three times for 20 min each, followed by two changes of propylene oxide. Tissue samples were then infiltrated with series of epoxy resin (25, 50, 75, 100%) for 24 h each and polymerized in the oven at 60°C for 48 h. The blocks were sectioned by an ultramicrotome (Ultracut E, Riechert-Jung), and sections of 80 nm were obtained and stained with uranyl acetate and lead citrate. Sections were observed using a Jeol JEM 1400 Plus transmission electron microscope and pictures were taken using a Gatan Orius CCD camera. EM micrographs were analyzed using computer-assisted ImageJ analysis software. To calculate g-ratio, myelinated axons (∼300, 100 axons per mouse, *n* = 3 per genotype) from EM images from corpus callosum were analyzed by dividing inner axonal diameter over the total axonal diameter.

### Open field test

The open field test was performed following the previously published protocol.^60^ Briefly, mice were placed in the corner of a 50 x 50 x 33 cm arena, and allowed to freely explore for 6 min. The center of the arena was defined as a 25 x 25 cm square in the middle of the arena. Velocity and time spent in the center and arena circumference were measured. Mice tested in the open field were recorded using a video camera connected to a computer having tracking software (Ethovision 12).

### Elevated plus maze test

The test apparatus consisted of two open arms (30 x 5 cm) bordered by a 1 cm high rim across from each other and perpendicular to two closed arms bordered by a rim of 16 cm, all elevated 75 cm from the floor. Mice were put into the maze and were allowed to explore it for 5 min. Duration of visits in both the open and closed arms were recorded.^60^

### Rotarod test

Each animal was placed on a rotating rod whose revolving speed increased from 5 rounds per min (rpm) to 40 rpm for 99 s. The test for each animal consisted of three trials separated by 20 min. Time to fall from device (latency) was recorded for each trial for each animal. If the animal did not fall from the device by 240 s from the beginning of the trial, the trial was terminated.^61^

### Image acquisition and analysis

Immunostained sections were imaged using a panoramic digital slide scanner or an Olympus FV1000 confocal laser scanning microscope or Nikon A1R+ confocal microscope. The acquired images were processed using the associated microscope software programs, namely CaseViewer, F-10-ASW viewer, and NIS elements respectively. Images were analyzed using ImageJ software. Images were analyzed while blinded to the genotype and the processing included the global changes of brightness and contrast.

### Statistical analysis

All graphs and statistical analyses was preformed using either Excel or GraphPad Prism 5. Results of the experiments were presented as mean ± SEM. The two-tailed unpaired Student’s t-test was used to test the statistical significance. Results were considered significant when the p <0.05, otherwise they were represented as ns (no significance). Data analysis was performed while blinded to the genotype. Sample size and p value is indicated in the figure legends.

---

**Supplementary Fig S1:**
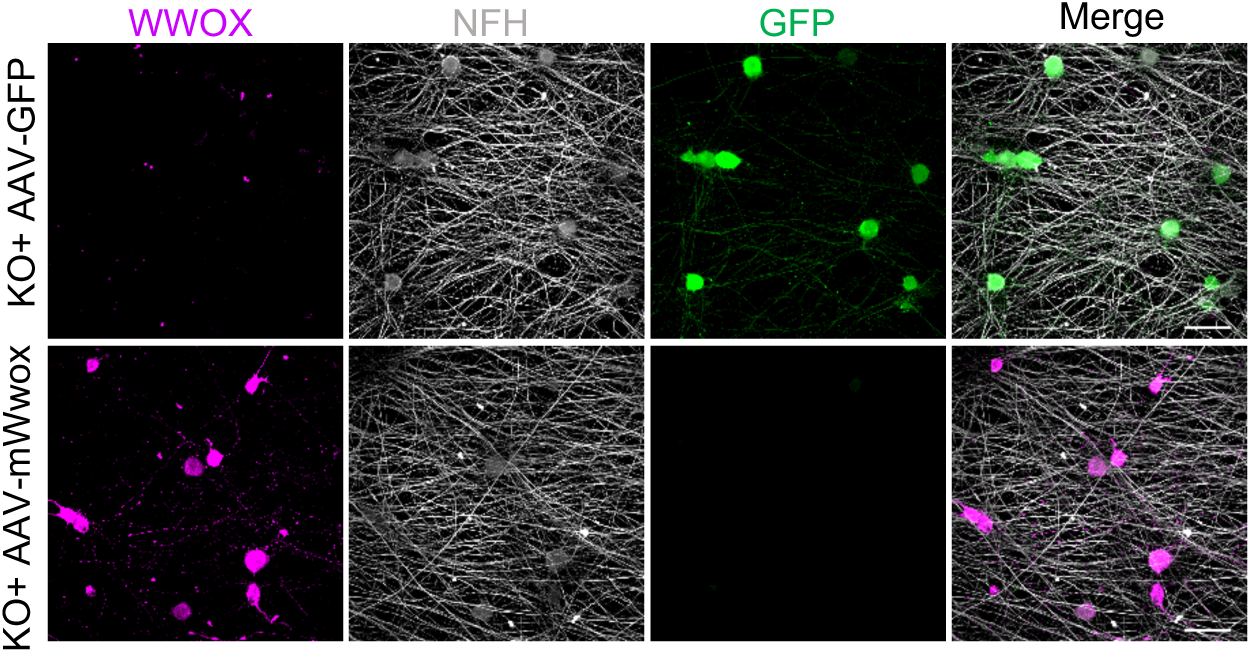
Validating the transgene expression in primary DRGs *ex vivo*. Cultured primary Wwox null DRGs were infected with AA9-hSynI-GFP or AA9-hSynI-mWwox. Immnuofluorescence showing the

**Supplementary Fig S2.**
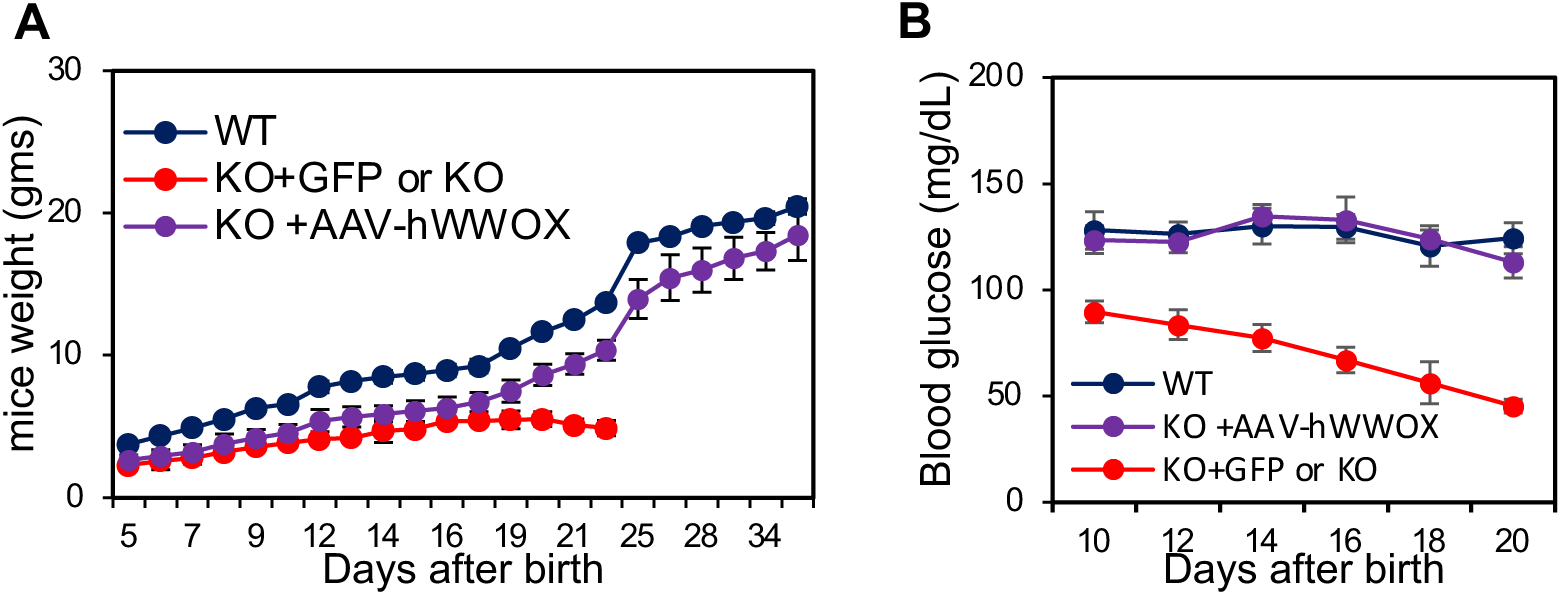
Effect of *AAV9-hSynI-hWWOX-IRES-GFP* on mice weight and blood glucose levels. at P17. (A) Graphs showing body weight (**A**) and blood glucose levels (**B**) of wild type mice (WT), *Wwox* null either injected with *AAV9-hSynI-GFP* (the control virus) or *AAV9-hSynI-hWWOX-IRES-GFP* virus at indicated days. Error bars represent ±SEM, n=4 mice per genotype.

**Supplementary Fig. S3.**
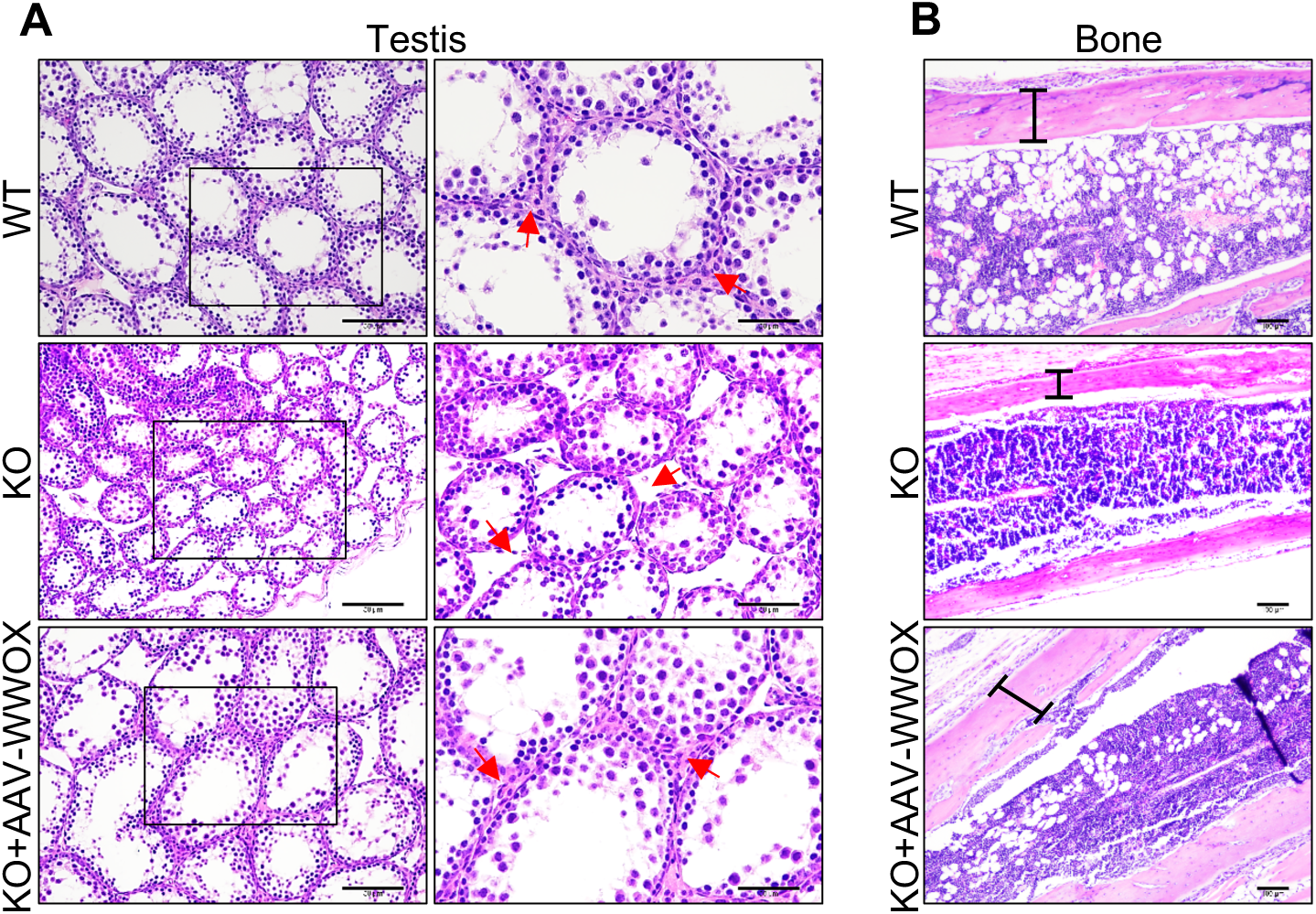
Neuronal WWOX restoration improves the development of testis and bone. **A**) Representative images showing the histology of testis from WT (n=4), KO (n=3), KO+AAV-WWOX (n=3) at P17. Arrows (red) indicate Leydig cells in WT and KO+AAV-WWOX and their absence in KO. Magnified area is shown with a box on the left panel. **B**) Representative histological (longitudinal section) images of bone (tibia) from WT, KO, KO+AAV-WWOX at P17. Bar represents bone width from WT (156±13.8µm, n=3), KO (38.5±6.8µm, n=3) and KO+AAV-WWOX (130±7.9µm, n=3). Bone width was measured from 3 different regions of tibia from each mice (n=3).

**Figure S4.**
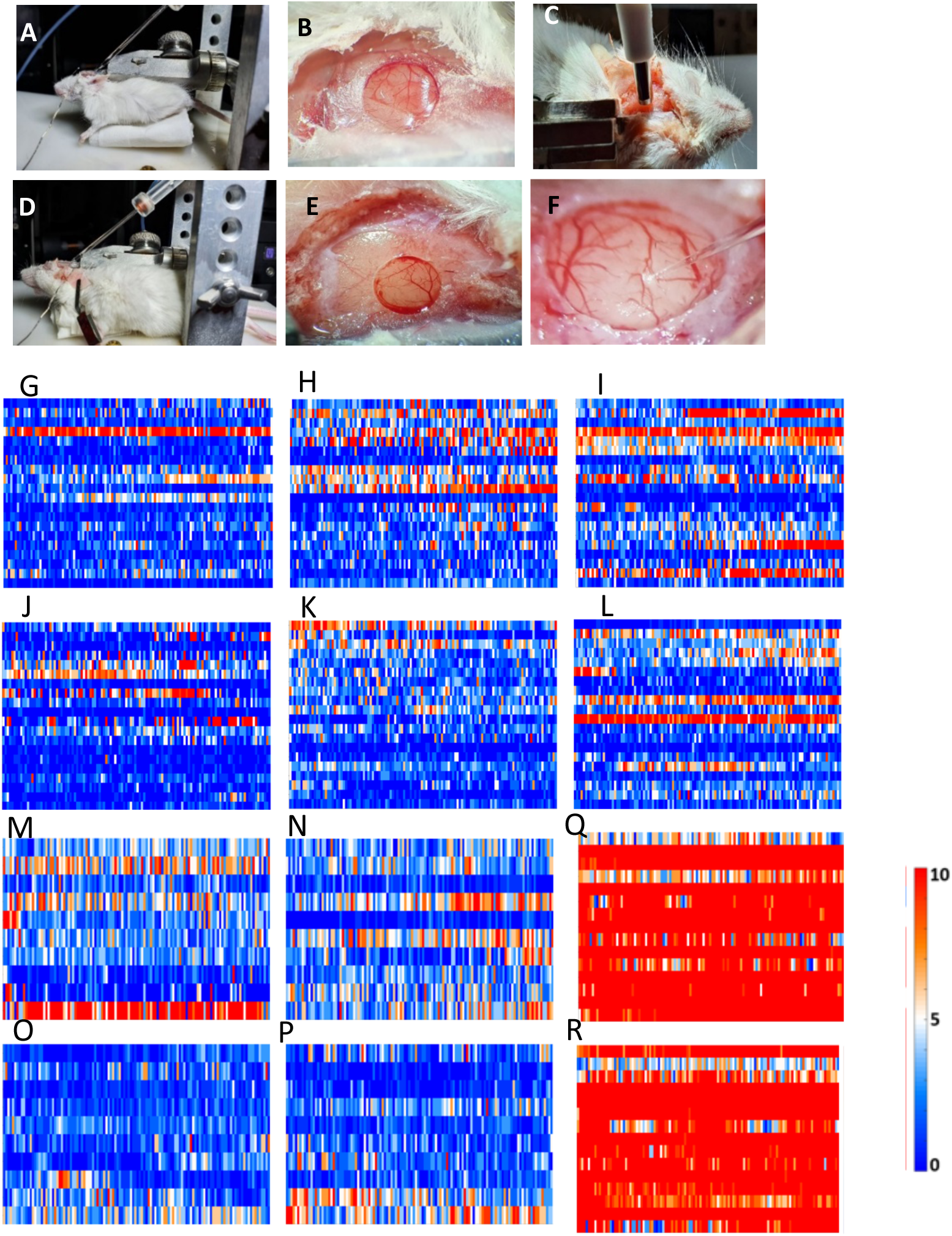
Surgical Procedures and Cell attached recordings of WT, KO (*Wwox* null) and AAV9*-mWwox*-treated mice. **a**) A view of a juvenile mouse (P20) placed on the stereotax with an electrode inserted into the brain. **b**) A 3-mm diameter craniotomy, positioned 1.6-2 mm posterior to the bregma and 4 mm lateral to the midline in a juvenile mouse. **c**) A view of an adult mouse (6 months) placed on the stereotax with an electrode inserted into the brain. **d**) A 3 mm diameter craniotomy, positioned 1.6-2 mm posterior to the bregma and 4 mm lateral to the midline in an adult mouse. **e**) A biopsy punch was used to cut through the skull. **f**) The electrodes were inserted at a 45 degrees and reached a depth of 200-300 µm. **g-i**) Raster plots of cell-attached recordings performed in adult WT mice (n=3) and a total of 60 recorded neurons (20 neurons from each animal). Each raster plot presents the results from one mouse. Every line within the raster plot presents one recorded neuron. The activity was recorded over 4 minutes and was binned at 2000 ms. The total amount of action potentials within the bin is color coded according to the colormap presented. **j-I**) Similar to g-I but recordings presented are from KO+A-Wwox mice (n=3). **m-n**) Raster plots of cell-attached recordings performed in WT pups (n=2) and a total of 20 recorded neurons (10 neurons from each pup). Each raster plot presents the results from one WT pup. Every line within the raster plot presents one recorded neuron. The activity was recorded over 4 minutes and was binned at 2000 ms. The total amount of action potentials within the bin is color coded according to the colormap presented o-p. Similar to m-n but recordings in KO+A-Wwox pups (n=2) (10 neurons in each of the pups). **q-r**) Similar to m-n but recordings in KO pups (n=2) (15 neurons in each of the pups). A clear hyperactivity was observed in the KO pups.

